# The MIR-NAT *MAPT-AS1* does not regulate Tau expression in human neurons

**DOI:** 10.1101/2023.01.27.525631

**Authors:** Rafaela Policarpo, Leen Wolfs, Saul Martínez-Montero, Lina Vandermeulen, Ines Royaux, Gert Van Peer, Pieter Mestdagh, Annerieke Sierksma, Bart De Strooper, Constantin d’Ydewalle

## Abstract

The *MAPT* gene encodes Tau protein, a member of the large family of microtubule-associated proteins. Tau forms large insoluble aggregates that are toxic to neurons in several neurological disorders and neurofibrillary Tau tangles represent a key pathological hallmark of Alzheimer’s disease (AD) and other ‘tauopathies’^1,2^. Several Tau-lowering strategies are being investigated as a potential treatment for AD but the mechanisms that regulate Tau expression at the transcriptional or translational level are not well understood^3,4^. Recently, Simone et *al*.^5^ reported the discovery of a new type of natural antisense transcripts called ‘MIR-NATs’ (referring to Mammalian-wide Interspersed Repeats – Natural Antisense Transcripts). Simone and colleagues used *MAPT-AS1*, a MIR-NAT associated to the *MAPT* gene as an archetype of this new class of long non-coding RNAs. According to Simone et *al*., *MAPT-AS1* represses Tau translation by competing for ribosomal RNA. We investigated the potential functions of *MAPT-AS1* in neurons using the same gain- and loss-of-function experiments as described in the original report and expanded our analysis with complementary approaches. Our data do not support a role for *MAPT-AS1* in regulating Tau expression in human neurons and urge to cautiously interpret the data provided by Simone et *al*.^5^.

## Results and Discussion

Natural antisense transcripts (NATs) are a particular class of long non-coding RNAs that can regulate expression of their overlapping protein-coding genes at the epigenetic, transcriptional or translational level^6^. Simone et *al*. identified *MAPT-AS1* transcripts as NATs that overlap with the *MAPT* gene and that contain embedded mammalian-wide interspersed repeat (MIR) motifs^5^. These motifs are complementary to sequences in the 5’ untranslated region (5’UTR) of the *MAPT* mRNA and were proposed to repress Tau translation by competing with ribosomal RNA pairing^5^. Simone et *al*. noted an enrichment of MIR-NATs in loci that harbor protein-coding genes associated to neurodegenerative disorders and/or that encode for intrinsically disordered proteins^5^.

To investigate whether long non-coding RNAs could regulate *MAPT* expression, we used a variety of bioinformatics approaches to identify putative NATs overlapping with the *MAPT* gene. We used the UCSC Genome Browser (https://genome.ucsc.edu/) and noted the presence of the *MAPT-AS1* locus on the opposite strand and partially overlapping with the first exon of *MAPT* (Fig. 1a). The genomic locus of *MAPT-AS1* spans approximately 52 kilobases; the mature reference transcript (RefSeq: NR_024559.1) is 840 nucleotides long and contains two exons that do not overlap with any of the *MAPT* exons (Fig. 1a). To confirm the identity of the mature transcript, we combined poly(A)+ RNA sequencing data from human brain reference RNA with FANTOM 5 capped analysis of gene expression (CAGE) sequencing data to assess read coverage in the *MAPT-AS1* locus (Fig. 1b), and more specifically at the annotated transcription start site (TSS) of *MAPT-AS1*. We also analyzed reads from 3’-end RNA-sequencing on human brain reference RNA (Fig. 1b,c). All three independent but complementary approaches confirmed expression of the reference sequence, with CAGE-seq and 3’end-seq read coverage coinciding with the annotated TSS and transcript end, respectively. To evaluate the presence of alternative *MAPT-AS1* isoforms, we performed *de novo* transcript assembly using poly(A)+ RNA-sequencing data from human brain reference RNA. One additional isoform was assembled with additional exons at the 3’ end but its expression levels in human brain were low and no supporting 3’-end coverage was observed for this isoform (data not shown). Our data thus supports the presence of the reference *MAPT-AS1* transcript of 840 nucleotides in length. A coding potential calculator^7^ and an open reading frame finder tool (https://www.ncbi.nlm.nih.gov/orffinder/) indicated that *MAPT-AS1* has no protein-coding potential and thus represents a putative NAT.

**Fig. 1.**
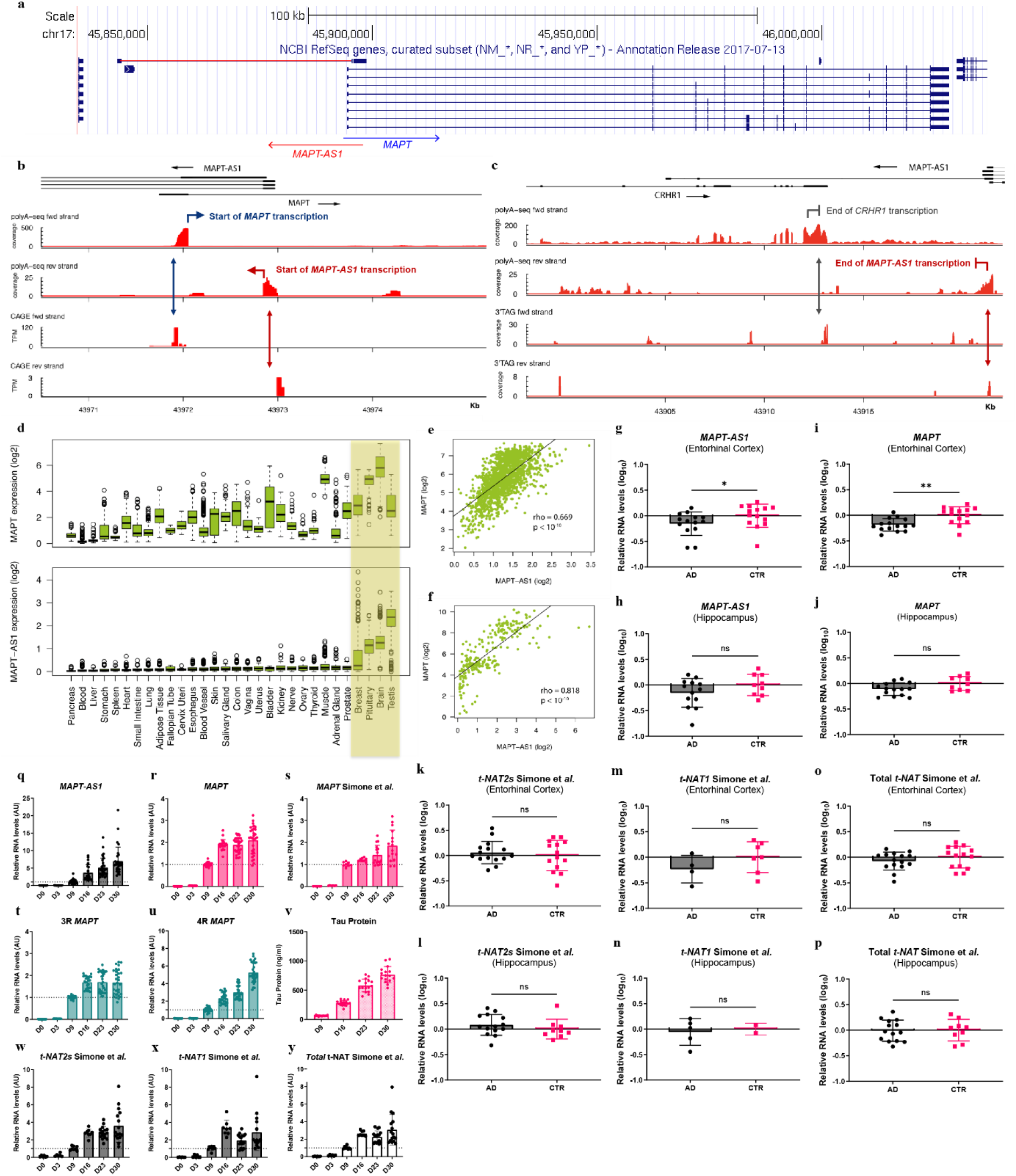
*MAPT-AS1* is a brain-enriched NAT that is dynamically co-expressed with the *MAPT* gene in human neurons. **a**, *MAPT-AS1* arises from the antisense strand of the *MAPT* gene in an intronic region downstream of its first exon and does not overlap with any *MAPT* exons. Source: https://genome.ucsc.edu/. **b, c**, Analysis of polyA+ RNA-seq data, CAGE-seq and 3’end-seq data in human brain reference RNA identified two potential *MAPT-AS1* isoforms. **d**, expression distribution of *MAPT* and *MAPT-AS1* across all tissue types in the GTex dataset; log_2_ TPM (transcripts per kilobase million) data is shown; tissues ordered based on *MAPT-AS1* expression levels. **e, f** Expression correlation between *MAPT* and *MAPT-AS1* in human brain samples from GTex (**e**) and FANTOM 5 CAGE datasets (**f**); log_2_ of TPM data was used; spearman’s *rho* values and p-values are shown. **g-p**, Expression levels of *MAPT-AS1* (**g, h**), *MAPT* mRNA (**i, j**) and *t-NAT* transcripts (**k-p**) in human brain samples from AD patients and Control individuals evaluated by RT-qPCR; *n* = 15 AD and 14 CTR for entorhinal cortex; *n* = 14 AD and 9 CTR for hippocampus; values scaled to Control (CTR) group; data are mean ± SD; Mann-Whitney test or unpaired t-test with df = 27 for entorhinal cortex and df = 21 for hippocampus, two-tailed p-value (*, p ≤ 0.05; **, p ≤ 0.001). **q-u**, **w-y**, Expression levels of *MAPT-AS1* (**q**), *MAPT* mRNA (**r-u**) and *t-NAT* transcripts (**w-y**) during differentiation of iNeurons at days 0, 3, 9, 16, 23 and 30 evaluated by RT-qPCR; *n* = 2 to 6 independent differentiations of iNeurons; values scaled to data from 9 days *in vitro*; data are mean ± SD. **v**, Tau protein levels (ng/mL) during neuronal differentiation of iNeurons at days 9, 16, 23 and 30 *in vitro* using a full-length Tau protein MSD assay; *n* = 2 independent differentiations of iNeurons; data are mean ± SD.

Simone et *al*. reported on three different transcript variants arising from the *MAPT-AS1* locus: *t-NAT1*, *t-NAT2l* and *t-NAT2s*^5^, where *t-NAT2s* represents the reference sequence. The relatively high number of reads spanning the first and last exon (52 reads) from *t-NAT2s* suggests that this transcript is the most abundant variant present in human brain. This is in line with our sequencing data. The very low number of exon-spanning reads that link the alternate exons of *t-NAT1* and *t-NAT2l* in the Simone et *al*.^5^ paper (1-7 reads) and the lack of good coverage of the alternate exons do not support robust expression of these transcript variants in human brain. Recent RNA sequencing data from the Telomere-to-Telomere (T2T) Consortium^8^ provide additional evidence that *t-NAT2l* is not detected (https://genome.ucsc.edu/). Despite differences in the identities of *MAPT-AS1* transcripts between our analyses and the work by Simone et *al*.^5^, we confirmed that the functionally relevant sequences including the MIR sequence and motifs 1, 2 and 3, described in the original report are present in the reference *MAPT-AS1* transcript (Extended data Fig. 1a).

To study the relationship between *MAPT-AS1* and *MAPT* expression, we used publicly available expression data^9,10^ and found, in line with Simone et *al*.^5^, that *MAPT-AS1* and *MAPT* are enriched and co-expressed in the brain (Fig. 1d-f). We further evaluated *MAPT-AS1* expression levels in two different regions of *post-mortem* brain samples from AD patients and age-matched controls. Consistent with publicly available data^9,10^, we detected both *MAPT-AS1* and *MAPT* in the entorhinal cortex and hippocampus isolated from controls and AD patients (Fig. 1g-j). *MAPT-AS1* and *MAPT* mRNA levels were reduced in the entorhinal cortex of AD brains as compared to controls (Fig 1g-j). These lower expression levels are likely the result of neuronal loss as suggested by the reduced expression of other neuronal genes in both brain regions (Extended Data Fig. 1c-f). Expression levels of general markers of neuroinflammation were increased in the same samples (Extended Data Fig. 1g-j). Our observations are in agreement with publicly available tools evaluating gene expression changes during AD progression (http://swaruplab.bio.uci.edu:3838/bulkRNA/)^11^ and with reports on *MAPT-AS1* expression in the brains of Parkinson’s disease patients^12,13^.

Then, we evaluated the expression of *t-NAT1*, *t-NAT2l* and *t-NAT2s* transcripts identified by Simone *et al*. in the same samples using the primer sequences designed by the original authors^5^ (Fig. 1k-p). In line with our RNA sequencing data, *t-NAT2s* was the main isoform present in human samples from both hippocampus and entorhinal cortex (Fig. 1k, l; Extended Data Fig. 1b). However, we found no expression of *t-NAT2l* in either patient group or brain region, and *t-NAT1* was expressed at very low levels and restricted to some individuals (Fig. 1m, n; Extended Data Fig. 1b).

To further investigate the cell type-specific expression of *MAPT-AS1*, we measured its levels in human induced pluripotent stem cell (iPSC)-derived microglia, astrocytes and neurons. Both *MAPT-AS1* and *MAPT* were particularly enriched in two models of human iPSC-derived neurons compared to astrocytes and microglia where no *MAPT-AS1* expression was detected (Extended Data Fig. 1k). To confirm that *MAPT-AS1* is predominantly expressed in neurons, we assessed its cellular distribution in the brain of AD patients and controls by *in situ* hybridization. *MAPT-AS1* localized predominantly to *MAPT*+ or *RBFOX3*+ neurons in control and AD brains (Extended Data Fig. 1l). Similar to Simone et. *al*^5^, we found a gradual increase in the levels of *MAPT-AS1*, *MAPT* transcripts and Tau protein during neuronal differentiation in a fast maturation inducible model of human iPSC-derived neurons called ‘iNeurons’ (Fig. 1q-y). This model has been used by other research groups to investigate Tau modulation approaches^14–16^ (Extended Data Fig. 1m). Although the differentiation protocols used to obtain human iPSC-derived neurons are different between our study and Simone *et*. al^5^, the expression dynamics of *MAPT-AS1* and *MAPT* are similar; both models can thus be used to investigate the functional role(s) of *MAPT-AS1*. In agreement with our previous human brain data, *MAPT-AS1/t-NAT2s* was the most abundant transcript in iNeurons (Fig. 1q, w; Extended Data Fig. 1b). We did not detect *t-NAT2l* expression in iNeurons, while *t-NAT1* was expressed at low levels in these cells using RT-qPCR assays developed by Simone and colleagues (Extended Data Fig. 1b).

The subcellular localization of long non-coding RNAs dictates largely its potential intracellular functions^17,18^. *In situ* hybridization on control and AD brains and RNA fractionation experiments confirmed that *MAPT-AS1* localized predominantly to the cytoplasmic fraction (Extended Data Fig. 1l, n, o). These observations are generally in line with the data reported by Simone et *al*.^5^ and suggest that *MAPT-AS1* may regulate Tau expression at the post-transcriptional or translational level in the cytoplasm of human neurons.

Simone et *al*. investigated the potential regulatory effects of the MIR-NAT *MAPT-AS1* by silencing the transcript variants in human neuronal clonal cells (SH-SY5Y cells) using small interfering RNAs (siRNAs) and in human iPSC-derived motor neurons by transduction with short hairpin (shRNA) constructs. Knockdown of *MAPT-AS1* did not alter *MAPT* mRNA levels but increased total Tau protein levels by ~50% relative to the control condition used. Conversely, Simone et *al*. reported a reduction of Tau protein levels by ~59-86% relative to the control condition when *MAPT-AS1* or a fusion construct containing only the critical motifs was expressed, while *MAPT-AS1* mutants not containing the MIR motifs lost their ability to inhibit Tau translation. Together, these data suggest that the MIR motifs in *MAPT-AS1* are essential to repress Tau expression during translation without affecting *MAPT* mRNA levels^5^.

To independently investigate whether *MAPT-AS1* regulates *MAPT* expression in human neurons, we utilized similar approaches but came to different conclusions. First, we utilized ASOs, hybrid DNA molecules 20 nucleotides in length, engineered to reduce *MAPT-AS1* levels via an RNase-H dependent mechanism. We screened a total of 42 ASOs tiling the mature reference *MAPT-AS1* transcript in a human neuroblastoma cell line (SK-N-MC) previously reported to express *MAPT-AS1*^12^ and identified several candidate lead ASOs that reduced *MAPT-AS1* levels by at least 50% (data not shown). Then, we treated these cells with serially diluted candidate lead *MAPT-AS1* ASOs and identified two lead ASOs (ASO-10 and ASO-16) that dose-dependently reduced *MAPT-AS1* (data not shown). We noted that the target sequence from the most potent *MAPT-AS1* ASO (ASO-10) is present in all *MAPT-AS1* isoforms described by Simone et *al*.^5^. Treatment with a positive control ASO targeting another long non-coding RNA (*MALAT1*) reduced its target in a dose-dependent manner without affecting *MAPT-AS1*, Tau mRNA or protein levels (Extended Data Fig. 2a, b). Treatment with a non-targeting (scrambled) ASO also did not change *MAPT-AS1* or Tau expression levels at any concentration tested (Extended Data Fig. 2c, d). Cells treated with the most potent *MAPT-AS1* ASO (ASO-10) at various concentrations showed unaltered *MAPT* mRNA and Tau protein levels (Extended Data Fig. 2e, f).

**Fig. 2.**
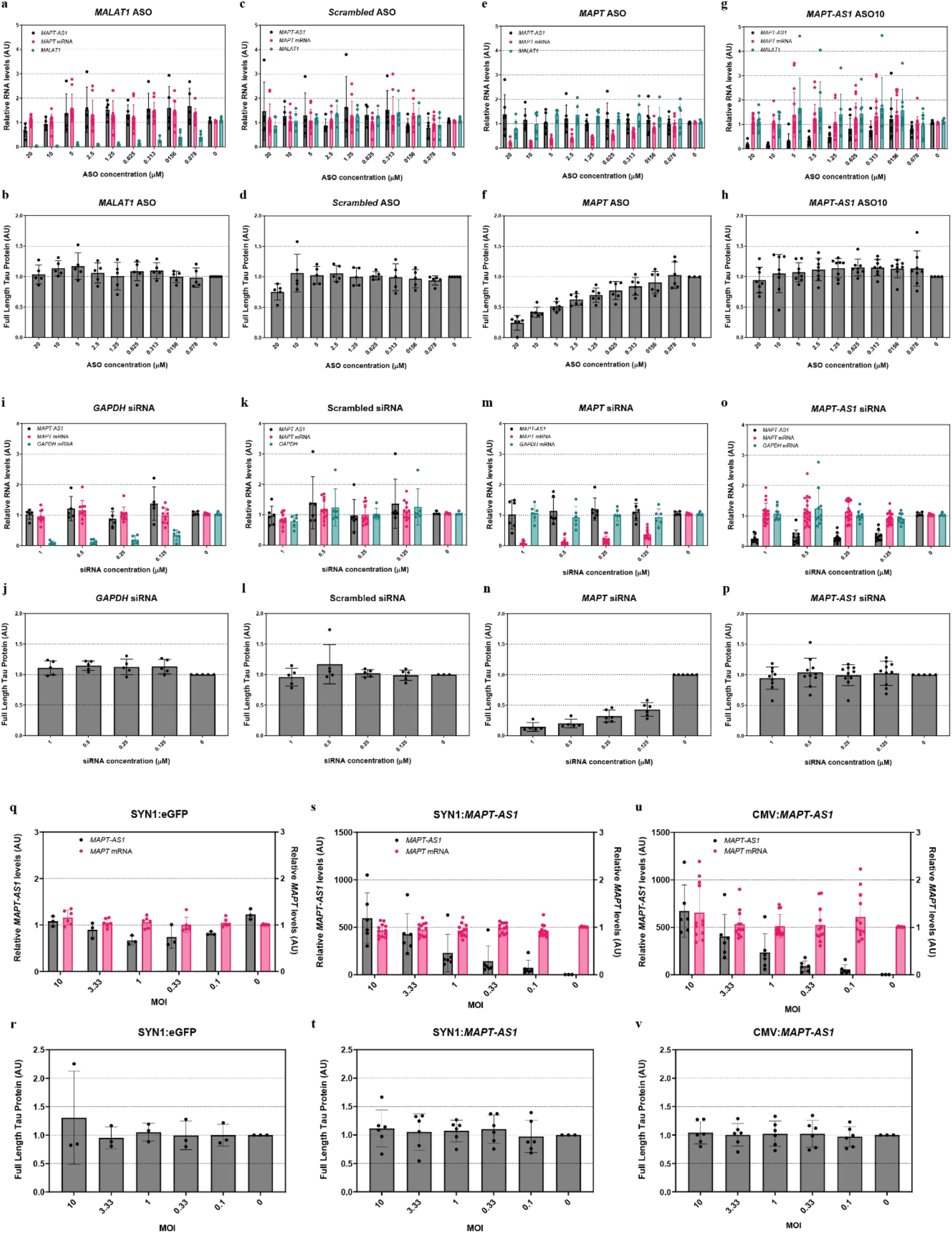
MAPT *-AS1* knockdown or overexpression does not affect Tau expression at the mRNA or protein level in iNeurons. **a-v**, iNeurons were treated with ASOs (**a-h**), siRNAs (**i-p**) or lentiviral constructs (**q-v**) at day 8 and harvested 10 days later at day 18 for RNA and Tau protein analysis. RNA expression levels were evaluated by RT-qPCR; values scaled to untreated condition. Tau protein levels were assessed using a full-length Tau protein MSD assay; values scaled to untreated condition (average set to 1); all data are mean ± SD. **a-h**, iNeurons treated with a *MALAT1* (**a, b**), a non-targeting (**c, d**), *MAPT* (**e, f**) or *MAPT-AS1* (**g, h**) ASOs; *n* = 3-4 independent experiments per ASO. **i-p**, iNeurons treated with a *GAPDH* (**i, j**), a non-targeting (**k, l**), *MAPT* (**m, n**) or *MAPT-AS1* (**o, p**) siRNAs; *n* = 3-5 independent experiments per siRNA. **a-h**, iNeurons treated with ASOs targeting *MALAT1* (**a, b**), a non-targeting ASO (**c, d**), *MAPT* (**e, f**) or *MAPT-AS1* (**g, h**); *n* = 3-4 independent experiments per ASO. **i-p**, iNeurons treated with siRNAs targeting *GAPDH* (**i, j**), a non-targeting siRNA (**k, l**), *MAPT* (**m, n**) or *MAPT-AS1* (**o, p**); *n* = 3-5 independent experiments per siRNA. **q-v**, iNeurons treated with lentiviral constructs overexpressing eGFP (**q, r**) or *MAPT-AS1* (**s-v**); *n* = 3 independent experiments per lentivirus. Overexpression of the transcripts was controlled by the synapsin-1 (SYN1) or the cytomegalovirus (CMV) promoter. MOI = multiplicity of infection.

To confirm our findings in differentiated human neurons, we treated iNeurons with several ASOs. A *MALAT1* ASO dose-dependently reduced *MALAT1* levels but did not alter Tau expression at the mRNA or protein level nor *MAPT-AS1* levels (Fig. 2a, b). Treatment with a scrambled ASO did not affect *MAPT-AS1* or Tau expression levels (Fig. 2c, d). We chose a long incubation time (10 days) to span the reported half-life (6-7 days) of Tau protein in human iPSC neurons^19^. We confirmed that this incubation time was sufficiently long to reduce Tau expression at the RNA and protein level in a dose-dependent manner using an ASO targeting *MAPT* mRNA (Fig. 2e, f). Reducing *MAPT* expression did not affect *MAPT-AS1* levels (Fig. 2e). Finally, treatment with the *MAPT-AS1* ASO dose-dependently reduced *MAPT-AS1* levels but did not change Tau mRNA or protein levels (Fig. 2g, h).

To rule out that our negative findings were attributed to the use of ASOs, we treated iNeurons neurons with a previously published custom-made siRNA targeting *MAPT-AS1*^12^. We confirmed that all *MAPT-AS1* transcripts annotated by Simone *et al*.^5^ contain the *MAPT-AS1* siRNA target sequence. A *GAPDH*-targeting positive control siRNA and a non-targeting siRNA did not affect *MAPT*-*AS1* nor Tau expression levels (Fig. 2i-l). Treatment with an siRNA targeting *MAPT* led to a dose-dependent decrease of *MAPT* mRNA and Tau protein levels (Fig. 2m, n). However, siRNA-mediated knockdown of *MAPT-AS1* did not result in altered *MAPT* expression at the RNA or protein level (Fig. 2o, p).

Finally, we also designed lentiviral constructs to overexpress the entire sequence of the reference *MAPT-AS1* transcript (including the MIR motifs) either using a neuronal-specific promoter (synapsin-1, SYN1) or a ubiquitously active promoter (cytomegalovirus-associated promoter, CMV). To ensure that potential effects on Tau protein could not be attributed to lentiviral toxicity, and in line with the claim that endogenous *MAPT-AS1* can regulate neuronal Tau protein levels in a sub-stoichiometric manner in motor neurons by Simone et *al*., we transduced iNeurons with up to 10 viral copies/cell. A positive control construct overexpressing eGFP did not change *MAPT-AS1* or *MAPT* expression levels relative to non-treated conditions (Fig. 2q,r). We also confirmed that both lentiviral constructs overexpressed *MAPT-AS1* in a dose-dependent manner in neurons (Fig. 2s,u). However, both *MAPT* mRNA and Tau protein levels remained unchanged (Fig. 2s-v). Similar results were obtained when we transiently overexpressed *MAPT-AS1* using the same constructs in SK-N-MC cells (data not shown).

To ascertain that the discrepancies between our observations and the findings summarized by Simone *et al*.^5^ were not caused by differences in cellular models and biochemical assays used, we sought to reproduce two key experiments from the original report. First, we evaluated the effect of reducing *MAPT-AS1* transcripts on Tau mRNA and protein levels in the same human neuroblastoma cell line (SH-SY5Y) using the same siRNA sequences and the same transfection procedures reported by Simone *et al*.^5^. We included a panel of positive and negative control siRNAs to confirm technical validity of our experiments. Our 2 positive control siRNAs targeting either *GAPDH* or *MAPT* mRNA did reduce their intended target mRNAs, whereas the negative control siRNA did not affect the levels of any RNA investigated in SH-SY5Y cells relative to non-treated cells (Fig. 3a-e; Extended Data Fig. 2q). In line with our previous observations but as opposed to Simone et *al*., we found no evidence for expression of *t-NAT2l* in SH-SY5Y cells (data not shown) and *t-NAT1* was not detected or expressed at variable levels in the same cellular model (Fig. 3d). Targeting *MAPT-AS1* using our siRNA resulted in a robust knockdown of *MAPT-AS1* and *t-NAT2s* transcripts (Fig. 3b,c). Treatment with siRNAs reported by Simone et *al*.^5^ resulted in variable knockdown levels of *MAPT-AS1* and its transcript variants (Fig 3a-e). Taken together, we were unable to confirm robust expression of 2 *MAPT-AS1* transcript variants in the same cellular model and using the same RT-qPCR assays as described by Simone et *al*.^5^. We were also not able to recapitulate a significant MAPT-AS1 or t-NAT transcripts knockdown in a neuronal cell line using the same siRNAs as originally described^5^.

**Fig. 3.**
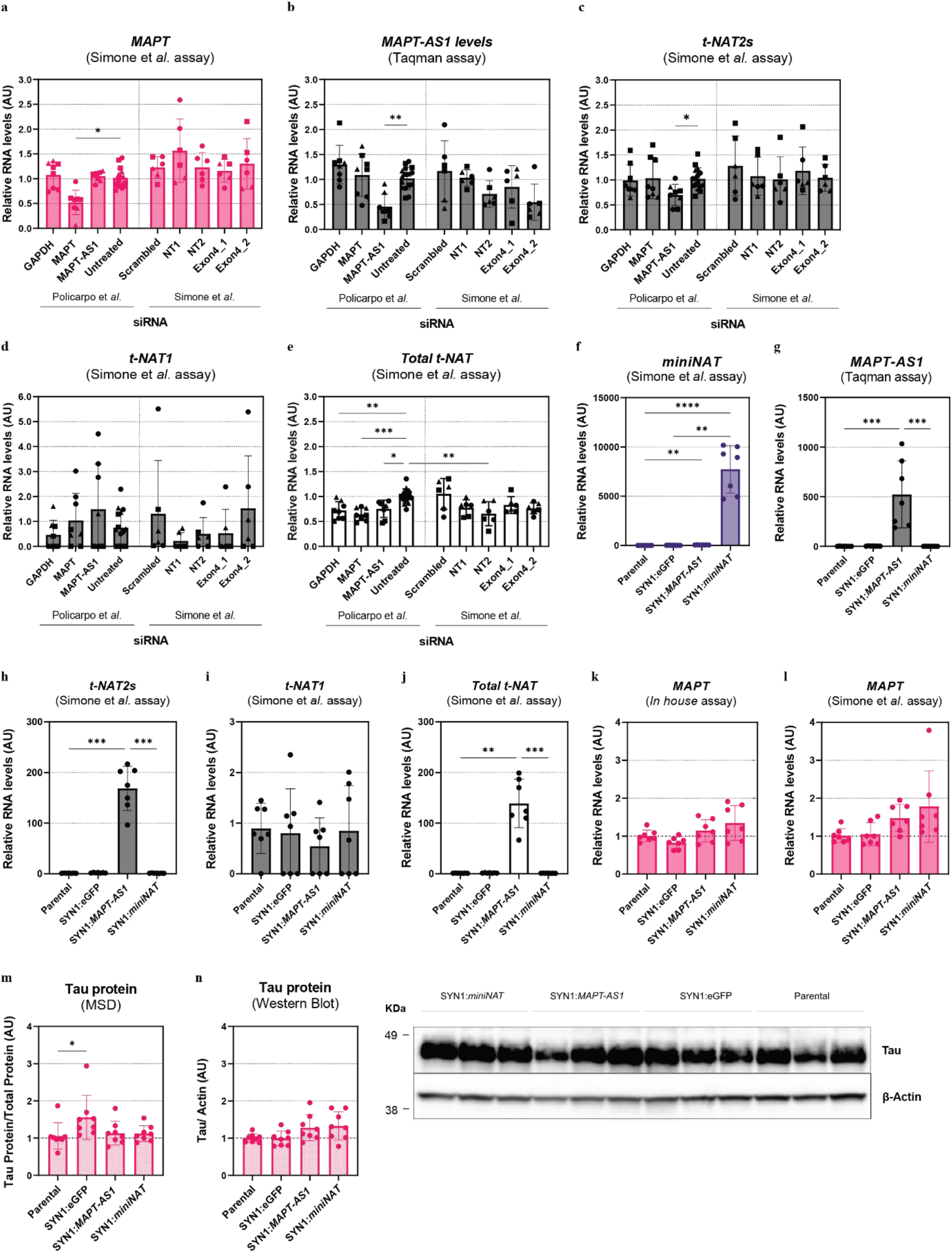
Transient siRNA-mediate reduction or stable overexpression of *MAPT-AS1* and *miniNAT* levels in SH-SY5Y cells. **a-e**, SH-SY5Y cells were treated with different siRNAs and harvested 48 hours later for RNA analysis. Results from siRNA sequences obtained from Simone et *al*.^5^ are shown on the right panel of each graph, as mentioned. Expression levels of *MAPT* mRNA (**a**), *MAPT-AS1* (**b**) and *t-NAT* transcripts (**c-e**) were evaluated by RT-qPCR; *n* = 3 independent experiments per siRNA; data from different experiments indicated as circles, squares or triangles, respectively; values scaled to non-treated condition; data are mean ± SD. Dunn’s multiple comparisons test (*, p ≤ 0.05; **, p ≤ 0.01; ***, p ≤ 0.001). **f-l**, SH-SY5Y cells were treated with lentiviral constructs at a multiplicity of infection (MOI) of 30 and cells stably expressing eGFP, *MAPT-AS1* or *miniNAT*, with exception for the parental line which remained non-treated. Cell lysates were harvested from 3 consecutive passages for RNA and Tau protein analysis. All data are mean ± SD. **f-j**, Expression levels of *miniNAT* **(f)**, *MAPT-AS1* (**g**) and *t-NAT* transcripts (**h-j**) were evaluated by RT-qPCR; *n* = 2-3 independent lysates per each condition and per passage; data from different experiments indicated as circles, squares or triangles, respectively; values scaled to parental line group; Dunn’s multiple comparisons test (*, p ≤ 0.05; **, p ≤ 0.01; ***, p ≤ 0.001; ****, p < 0.0001). **k, l**, *MAPT* mRNA expression levels in SH-SY5Y stable cell lines were evaluated by RT-qPCR using two independent primer sets; *n* = 2-3 SH-SY5Y lysates per passage; values scaled to parental line group; Dunn’s multiple comparisons test. **m, n**, Tau protein levels from SH-SY5Y lines were assessed using a full-length Tau protein MSD assay and normalized to total protein (**m**) or Western Blot analysis and normalized to β-actin levels (**n**); *n* = 2-3 independent lysates per each condition and per passage; values scaled to parental line group; One-way ANOVA with Holm-Šídák’s multiple comparisons test (*, p ≤ 0.05).

Secondly, we generated the same SH-SY5Y cells stably overexpressing *miniNAT* as the minimally required sequence to reduce Tau protein translation as reported by Simone et *al*.^5^. We included SH-SY5Y cells stably overexpressing eGFP and *MAPT-AS1* as negative and positive control, respectively. After selection, stably expressing cells showed robust and homogenous eGFP expression confirming similar transduction efficiencies across the different constructs (Extended Data Fig. 2g). We noticed that *MAPT-AS1*, *t-NAT* transcripts and Tau mRNA and protein levels varied with number of passages in SH-SY5Y cells (Extended Data Fig. 2h-p). Therefore, to ensure that any potential changes on Tau expression levels are dependent on *MAPT-AS1*, we analyzed both stable and parental SH-SY5Y cell lines at the same passage numbers. We confirmed stable overexpression of both *MAPT-AS1* (also representing its endogenously expressed transcript variants) and *miniNAT* over consecutive passages in SH-SY5Y cells (Fig. 3f-j). However, stable overexpression of *miniNAT* or *MAPT-AS1* did not show significant changes on *MAPT* mRNA levels (Fig. 3k,l) and did not result in decreased Tau protein levels compared to non-treated conditions when evaluated by the quantitative and sensitive MSD assay or using the same antibody for Western blot analysis as reported by Simone et *al*.^5^ (Fig. 3m,n). Finally, no changes on Tau expression levels were observed in iNeurons transduced with a virus overexpressing the *miniNAT* reported by Simone et *al*.^5^ (Extended Data Fig. 3a-f). The robust expression of eGFP signal in neurons after treatment with the SYN1:eGFP construct confirmed a high efficiency of transduction in this model (Extended Data Fig. 3g). Taken together, we were unable to confirm that overexpression of *miniNAT* reduces Tau protein translation in human neuronal cells as described by Simone et *al*.^5^.

## Conclusions

In summary, we identified a NAT associated to the *MAPT* gene that shows several similarities with the *MAPT*-associated MIR-NAT reported by Simone et *al*.^5^. First, *MAPT-AS1* is enriched in the human brain and its expression is reduced in the entorhinal cortex of AD brains, likely due to neuronal loss during disease progression. Second, *MAPT-AS1* localizes predominantly in the cytoplasm of neurons and its expression increases with neuronal maturation. However, and in sharp contrast to the observations made by Simone et *al*.^5^, either increasing or reducing *MAPT-AS1* levels had no effect on *MAPT* transcription or splicing (data not shown) nor on Tau translation in our models. Importantly, when using the same cellular models, the same biochemical tools and readouts, we were unable to recapitulate key experiments reported by Simone et *al*.^5^. Thus, we are unable to validate the biological relevance of the MIR-NAT *MAPT-AS1* as a negative regulator of Tau protein expression in human neurons.

Our sequencing methods imply that *MAPT-AS1* contains structural features (including a poly(A) tail and a 5’ cap consistent with other functional lncRNAs. Its cytoplasmic localization suggests that *MAPT-AS1* may play a role in post-transcriptional/translational mechanisms. Consistent with its cellular localization, we found no biologically significant changes in the transcriptome of human iPSC-derived neurons in which *MAPT-AS1* levels were either lowered by ASO treatment or increased by lentiviral overexpression (data not shown). Additional studies will be required to unravel the potential function(s) of *MAPT-AS1*. Several pharmaceutical companies are evaluating safety and efficacy of Tau-targeting therapeutic modalities in clinical trials (Reviewed in ^1,4^). Thus, we encourage the scientific community to continue exploring the mechanisms that regulate Tau expression in order to support the Tau-targeting strategies currently in clinical trials. Ultimately, this will pave the way for the development of novel therapeutics for AD and other tauopathies.

## Acknowledgments

We thank the London Neurodegenerative Diseases Brain Banks, the Netherlands Brain Bank, all the individuals and their families for providing human brain tissue samples. We would like to acknowledge Dr. Wei-Ting Chen for helping with the human brain cryosections to perform multiplex fluorescent *in situ* hybridization. We thank Dr. Lujia Zhou and Dr. Alfredo Cabrera for providing us with lysates from human iPSC derived astrocytes and Dual-Smad neurons. RP holds a doctoral student fellowship from VLAIO, the Flemish Agency for Innovation and Entrepeneurship (Vlaams Agentschap Innoveren & Ondernemen). This work was supported by VLAIO (R&D grants HBC.2018.2290 and HBC.2020.3236), and European Research Council (ERC) grant CELLPHASE_AD834682 (EU), FWO, KU Leuven, VIB, Stichting Alzheimer Onderzoek, Belgium (SAO), the UCB grant from the Elisabeth Foundation, a Methusalem grant from KU Leuven and the Flemish Government and Dementia Research Institute - MRC (UK). BDS is the Bax-Vanluffelen Chair for Alzheimer’s Disease and is supported by the Opening the Future campaign and Mission Lucidity of KUL, Leuven University.

## Author Contributions

RP and CdY were responsible for the conception and design of the project and the article with contributions from AS and BDS. IR designed and selected the lentiviral constructs and siRNAs used in this study. SMM designed and synthesized the *MAPT-AS1* ASOs. RP performed all experiments with human brain samples, SK-N-MC, SH-SY5Y and NGN2-inducible neurons. LV and CdY contributed to SK-N-MC and NGN2-inducible neurons experiments. RP and LW performed multiplex fluorescent *in situ* hybridization experiments in human brain samples. GVP and PM performed all bioinformatic analysis. RP and CdY analysed the data and interpreted results with contributions from AS, LV and BDS. All authors reviewed and approved the manuscript.

## Data Availability

All data that support the findings of this study are available from the corresponding author CdY upon request.

---

**Extended Data Fig. 1.**
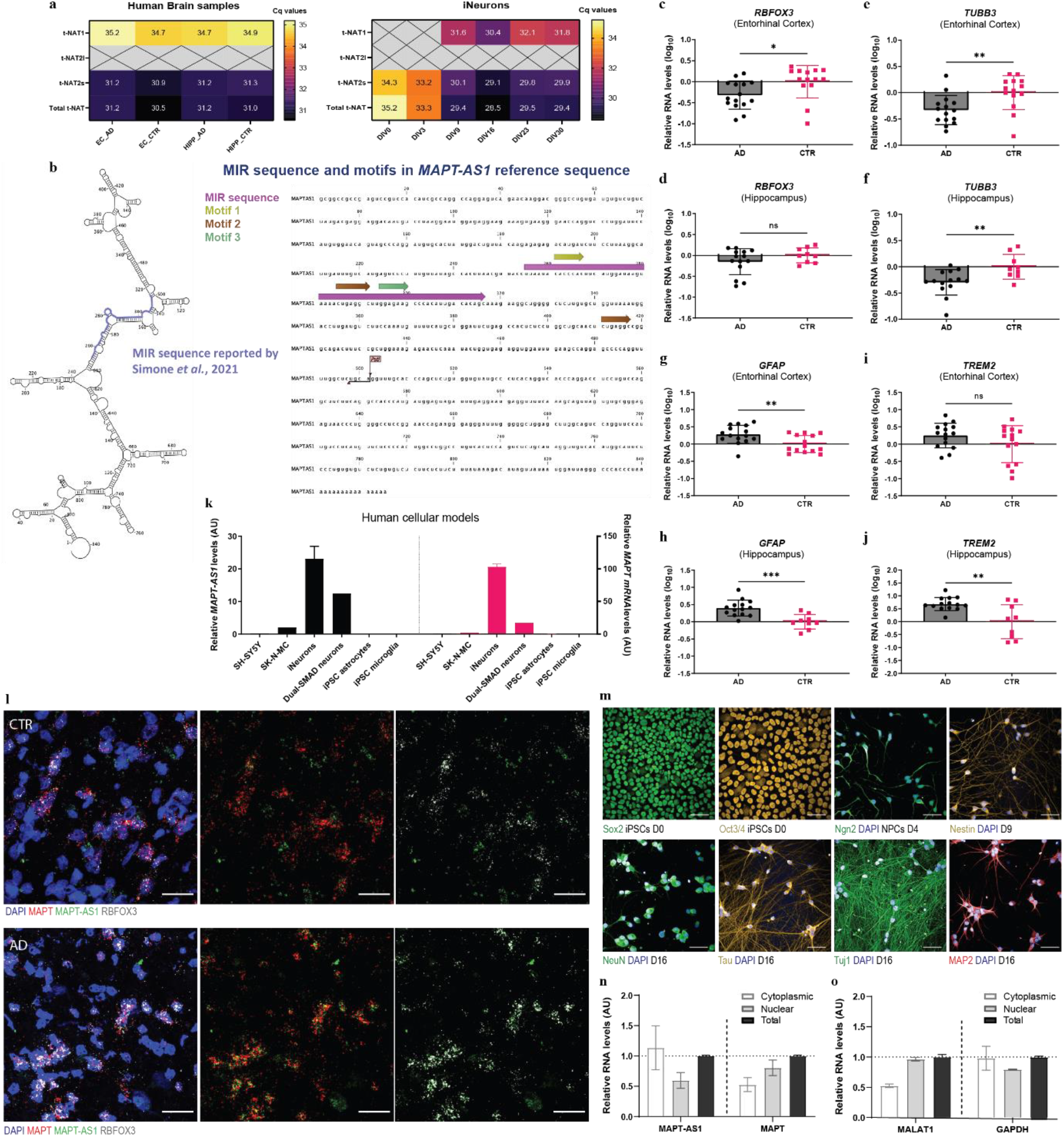
MAPT*-AS1* localizes predominantly to the cytoplasm of human neurons. **a**, Scheme of the annotation of MIR sequences reported by Simone et *al*.^5^ in the human *MAPT-AS1* reference transcript. Structure predictions were performed using CLC Main Workbench v.8.1.3 (Qiagen). **b**, Cq values from t-NAT transcripts in human brain (left panel) and in iNeurons (right panel) samples were obtained with RT-qPCR. Crossed samples correspond to no amplification. Data are mean. **c-j**, mRNA expression levels of neuronal markers *RBFOX3* (**c, d**) and *TUBB3* (**e, f**); and reactive astrocytic and microglial markers *GFAP* (**g, h**) and *TREM2* (**i, j**), respectively, in human brain samples from AD patients and Control individuals evaluated by RT-qPCR; *n* = 15 AD and 14 CTR for entorhinal cortex; *n* = 14 AD and 9 CTR for hippocampus; values scaled to Control (CTR) group; data are mean ± SD; Mann-Whitney test or unpaired t-test with df = 27 for entorhinal cortex and df = 21 for hippocampus, two-tailed p-value (*, p ≤ 0.05; **, p ≤ 0.001). **k**, Expression levels of *MAPT-AS1* and *MAPT* mRNA in SH-SY5Y and SK-N-MC cells, and human iPSC derived models were evaluated by RT-qPCR; *n* = 1-2 samples per cell type; astrocytes and microglia analysed at day 126 and day 14, respectively; iNeurons or human iPSCs differentiated to cortical neurons using the Dual-SMAD at days 30 and 28 *in vitro*, respectively; values scaled to average; data are mean (± SD, when applicable). **l**, *In situ* hybridization showing *MAPT* (red), *MAPT-AS1* (green) and *RBFOX3* (white) expression in *post-mortem* brains from CTR and AD patients. Representative images from 1 CTR and 1 AD patient. Scale bar 30μm. **m**, Human iPSCs show expression of stem cell markers Sox2 and Oct3/4 at day 0, and NGN2 after treatment with doxycycline for 4 days. Immature neurons at day 9 displayed a neuron-like morphology accompanied by expression of Nestin. Neuronal maturation markers MAP2, Tau, TUJ1 and NeuN are expressed at day 16. DAPI staining was used to detect cell nuclei. Scale bar 50μm; *n* = 4 independent differentiations of iNeurons. **n, o**, Expression levels of *MAPT-AS1*, *MAPT, MALAT1* (nuclear control) and *GAPDH* (cytoplasmic control) were evaluated by RT-qPCR in sub-cellular fractions of NGN2-inducible neurons at DIV23; *n* = 2 lysates from 1 differentiation of iNeurons; values scaled to total fraction; data not normalized to reference genes and shown as mean ± SD.

**Extended Data Fig. 2.**
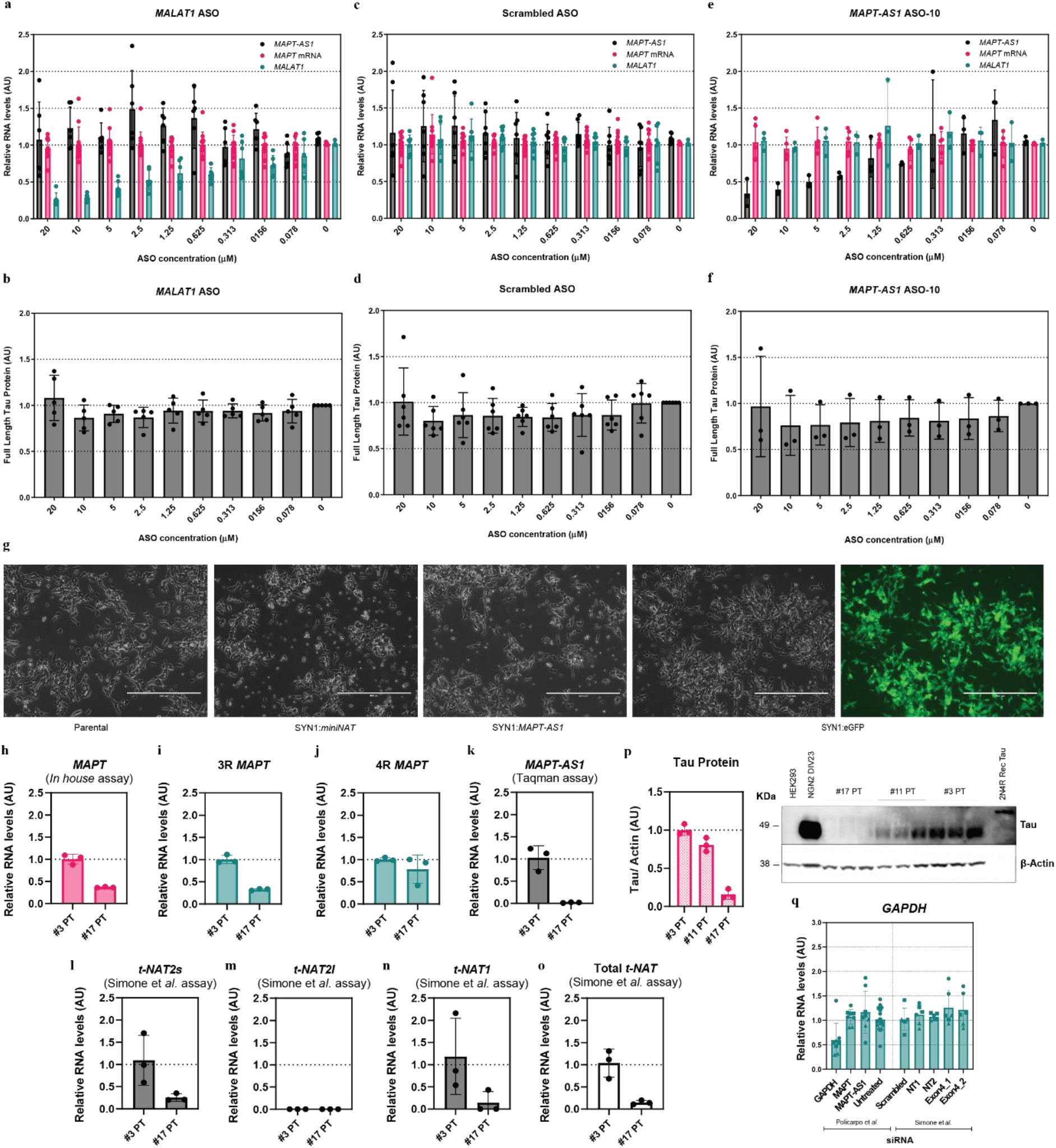
Modulation of *MAPT-AS1* levels in SK-N-MC and SH-SY5Y cells. **a-f**, SK-N-MC cells were treated with a *MALAT1* ASO (**a, b**), a non-targeting ASO (**c, d**), or the lead *MAPT-AS1* ASO (**e, f**) and harvested after 72 hours for RNA and Tau protein analysis. *n* = 3-4 independent experiments per ASO; RNA expression levels were evaluated by RT-qPCR; values scaled to untreated condition. Tau protein levels were assessed using a full-length Tau protein MSD assay; values scaled to untreated condition (average set to 1); all data are mean ± SD; **g**, eGFP expression in SH-SY5Y cells was confirmed after treatment with a SYN1:eGFP construct at a multiplicity of infection (MOI) of 30. Representative images 72 hours after puromycin treatment. Scale bar 400μm. **h-o**, Expression levels of *MAPT* (**h-j**) or *MAPT-AS1* and t-NAT (**k-o**) transcripts in SH-SY5Y cells at passage 3 or 17 post-thawing (#3 or #17 PT, respectively). RNA expression levels were evaluated by RT-qPCR; *n* = 3 SH-SY5Y lysates per passage; values scaled to lowest passage number (#3 PT); data are mean ± SD. **p**, Tau protein levels in SH-SY5Y cells at passage 3, 11 or 17 post-thawing (#3, #11 or #17 PT, respectively) were assessed using Western Blot analysis; *n* = 3 SH-SY5Y lysates per passage; Tau protein levels were normalized to β-actin levels and scaled to the lowest passage number (#3 PT; average set to 1); data are mean ± SD. **q**, *GAPDH* mRNA expression levels in SH-SY5Y cells treated with different siRNAs were evaluated by RT-qPCR; *n* = 3 independent experiments per siRNA; data from different experiments shown in circles, squares and triangles, respectively; values scaled to non-treated condition; data are mean ± SD.

**Extended Data Fig. 3.**
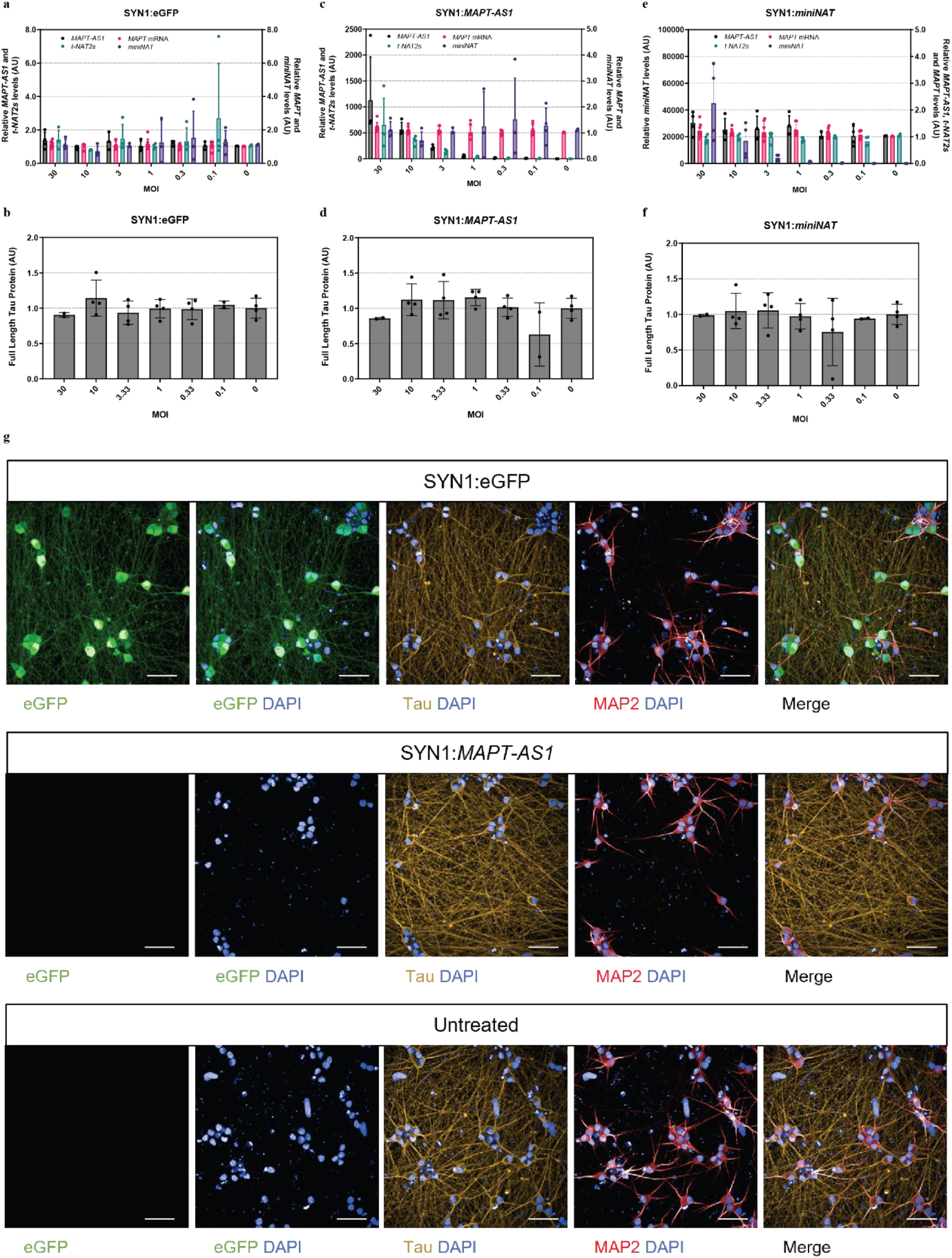
*miniNAT* overexpression does not affect Tau expression at the mRNA or protein level in iNeurons. iNeurons were treated with lentiviral constructs overexpressing eGFP (**a, b**), *MAPT-AS1* (**c, d**) or *miniNAT* constructs (**e, f**) at day 8 and harvested 10 days later at day 18 for RNA and Tau protein analysis; *n* = 3 independent experiments per lentivirus. RNA expression levels were evaluated by RT-qPCR; values scaled to untreated condition. Tau protein levels were assessed using a full-length Tau protein MSD assay; values scaled to untreated condition (average set to 1); all data are mean ± SD. **g**, eGFP expression in NGN2-inducible neurons was confirmed after treatment with a SYN1:eGFP construct at an MOI of 30. Representative images 10 days after treatment. Scale bar 30μm.

