## Supplementary material for "The MIR-NAT *MAPT-AS1* does not regulate Tau expression in human neurons": Methods

### Bioinformatic Analysis

To identify different *MAPT-AS1* isoforms, we performed a transcript-assembly analysis using the StringTie algorithm (v1.3.3b)[1] on internally generated stranded poly(A)+ RNA-sequencing data from human brain reference RNA (ThermoFisher). A library was generated using the TruSeq stranded mRNA library prep kit (Illumina) according to the manufacturer's instructions and sequenced on a NextSeq 500 (Illumina). Reads were mapped to the human reference genome (hg38) using TopHat (v2.1.0)[2]. Transcript assembly was guided by the *MAPT-AS1* reference annotation (Ensembl Release 90). 3'end-sequencing data were generated for from human brain reference RNA (ThermoFisher) using the QuantSeq library prep procedure (Lexogen) according to the manufacturer's instructions and sequenced on a NextSeq 500 (Illumina). Reads were mapped to the human reference genome (hg38) using TopHat.

Expression correlation between *MAPT* and *MAPT-AS1* was evaluated in the GTex RNA sequencing[3] and FANTOM 5 CAGE sequencing[4] datasets. Analyses were performed at the gene level. All analyses were performed using the R statistical programming language.

### Human Neuroblastoma Cell Lines (SK-N-MC and SH-SY5Y)

SK-N-MC (HTB-10) and SH-SY5Y (CRL-2266) cells were obtained from the American Type Culture Collection (ATCC). SK-N-MC cells were maintained in Minimum Essential Medium (MEM; Gibco) supplemented with 10% (v/v) heat-inactivated fetal bovine serum (HI-FBS, Biowest), 1 mM sodium pyruvate (Gibco), 1.5 g/L sodium bicarbonate, 0.1 mM non-essential amino acids (NEEA; Gibco) and 50 µg/ml gentamycin (Gibco). SH-SY5Y cells were maintained in Dulbecco's Modified Eagle's Medium/Nutrient Mixture F-12 Ham (DMEM/F12; Sigma) supplemented with 10% (v/v) HI-FBS (Biowest), 0.1 mM NEEA (Gibco) and 50 µg/ml gentamycin (Gibco). Cells were kept in a humidified incubator at 37 °C and 5% $CO_2$.

### SH-SY5Y stable cell lines

For establishing the stable cell lines (SYN1:MAPT-AS1; SYN1:miniNAT; SYN1:eGFP), SH-SY5Y cells were seeded in 96-well µclear plates (Greiner Bio-One) at a density of 40,000 cells per well and 24 hours later transduced with 3 custom designed lentiviral constructs, overexpressing, under the neuronal synapsin 1 (SYN1) promoter, the following sequences: 1) the entire cDNA sequence of *MAPT-AS1* (NR_024559.1) except for the PolyA-tail sequence (SYN1:*MAPT-AS1*), 2) the *miniNAT* sequence provided by Simone et *al.*[5] (SYN1:*miniNAT*) and 3) a control lentiviral construct overexpressing eGFP (SYN1:eGFP) at a multiplicity of infection (MOI) ratio of 30. MOI = Virus Titer / Number of Cells. A total of 6 wells was transduced per lentiviral construct; a total of 18 wells remained untreated (parental line). The day after transduction, media containing lentiviral particles was replaced by fresh media in each well. On the next day, fresh media containing 1 µg/mL puromycin was added to lentiviral-treated conditions to start selection of stable cell colonies. Cells were cultured for another 5 days with regular media replacement with fresh puromycin containing media until resistant colonies could be identified. Then, cells from 3 independent wells per condition (done twice to expand independent colonies) were split together and transferred into a total of 6 wells per condition in 24-well plates (Falcon). Cells were cultured for another 10 days with regular media replacement with fresh puromycin containing media before being transferred to t25 flasks (two flasks per condition to expand independent colonies). Cells were then plated in 6-well plates at a density of 300,000 cells/well over 3 consecutive passages. Independent lysates from each stable and parental SH-SY5Y cell lines were collected through 3 consecutive passages (#4, #5 and #6 post-thawing, PT). At least 2 wells per condition and per passage were collected for RNA and protein analysis.

Vector construction and lentiviral production done at VectorBuilder. Information on lentiviral constructs is available at Supplementary Table S1. The sequences and vector maps of the lentiviral constructs can be provided upon request by e-mail to corresponding author.

### Differentiation of Human iPSC-derived microglia

Human induced pluripotent stem cells (hiPSC)-derived microglia were obtained from the ApoEε3/3 hiPSC line (UKBi011-A-3), an isogenic mutant of the ApoEε4/4 genotype parental line (UKBiO011-A)[6]. After thawing the hiPSCs on Matrigel (Corning) coated plates in mTeRS1 medium (Stem Cell Technologies) containing 10 µM ROCK inhibitor (ROCKi, Sigma), the cells were cultured, and the medium changed daily with fresh mTeSR1 medium without ROCKi. When confluency was reached, the cells were passaged with EDTA (Gibco), and differentiated into embryonic bodies (EBs), macrophage precursor factories (MPFs) and microglia using a method described by Sally Cowley *et al.*[7,8]. Briefly, hiPSCs were dissociated into single cell suspension and EBs were produced in Aggrewell plates (Stem Cell Technologies) by addition of 0.05 µg/mL BMP-4 (Invitrogen), 0.05 µg/mL VEGF (PeproTech) and 0.02 µg/mL SCF Miltenyi into mTeSR1 medium. After three days of daily medium changes, the EBs were transferred to 6-well factory plates (ThermoFisher) to create MPFs by placing 10-20 EBs per well in X-VIVO15 medium (Lonza) containing 10 U/mL Penicillin-Streptomycin (Gibco), 1% (v/v) GlutaMAX (Gibco), 50 µM 2-Mercaptoethanol (Gibco), 0.1 µg/mL M-CSF (Gibco) and 0.025 µg/mL IL-3 (Gibco). The factories were cultured for about 6-8 weeks with weekly medium changes before the suspension cells produced by the MPFs could be collected for final differentiation into functional and mature microglia. Microglia were cultured at a density of 15,000 cells per well in uncoated 96-well µclear plates (Greiner Bio-One), in Advanced DMEM/F12 medium (Life technologies) containing 10 U/mL Penicillin-Streptomycin (Gibco), GlutaMAX, 50 µM 2-Mercaptoethanol (Gibco), 0.01 µg/mL GM-CSF (Life Technologies) and 0.1 µg/mL IL-34 (Peprotech) for 14 days, with half medium changes every 2-3 days.

### Differentiation of Human iPSC-derived cortical neurons (Dual-Smad) and Human iPSC-derived astrocytes

hiPSCs were differentiated into cortical neuronal progenitor cells (NPCs) by Axol Biosciences (Cambridgeshire, UK), as previously described[9]. Cells were thawed in N2B27 neuronal differentiation medium [1:1 mixture of Neurobasal medium (Gibco) with DMEM:F12 Glutamax (Gibco) with 1% (v/v) B27 supplement (Gibco), 0.5% (v/v) GlutaMAX (Gibco), 0.5% (v/v) N2 supplement (Gibco), 2.5 µg/mL Insulin solution, 25 µM 2-Mercaptoethanol (Gibco), 0.5% (v/v) NEEA (Gibco), 0.5 mM sodium pyruvate (Gibco) and 10U/mL Penicillin-Streptomycin (Gibco)], supplemented with 20 ng/mL of human recombinant basic fibroblast growth factor (bFGF, Life Technologies).

For neuronal differentiation, cells were dissociated three days after thawing (experimental day 0) using Accutase (Sigma) and seeded on 24-well culture plates (Corning) coated overnight with undiluted Poly-L-ornithine solution (PLO, Sigma) followed by 10 µg/mL of mouse laminin (Sigma), at a density of 125,000 cells per well in neuronal differentiation medium supplemented with 10 µM ROCKi. From day 1 onwards, media changes were done once a week by replacing half of the medium with N2B27 neuronal differentiation media supplemented with 20 ng/mL BDNF (R&D Systems), 20 ng/mL GDNF (R&D Systems), 1 mM db-cAMP (Sigma) and 200 µM L-AA2P (Sigma) until sampling at day 28.

For differentiation of NPCs into astrocytes, cells were dissociated with Accutase (Sigma) three days after thawing (experimental day 3) and seeded on 6-well culture plates (Corning) coated overnight with undiluted PLO (Sigma) followed by 10 µg/mL of mouse laminin (Sigma) at a density of 750,000 cells/well in Astrocyte induction media [N2B27 supplemented with 20 ng/mL of human recombinant CNTF (R&D Systems)]. Between day 12 and 19, medium was switched to Astrocyte Differentiation medium [MEM medium supplemented with 0.6% (v/v) D-glucose (Milipore) and 10% (v/v) HI-FBS)] and changed once per week. Astrocytes were cultured for 7-8 weeks and dissociated/sub-plated with Accutase (Sigma) several times until sampling at day 126.

### Differentiation of Human iPSC-derived NGN2-inducible Neurons (iNeurons)

BIONi010-C-13 gene edited hiPSC cell line was obtained from the European Bank for induced pluripotent Stem Cells (https://cells.ebisc.org/BIONi010-C-13)[10]. Cells were first genotyped for their *MAPT* haplotype and confirmed to be H1/H1 homozygous. To generate NPCs, hiPSCs were thawed and seeded on Matrigel (Corning) coated 6-well plates (Corning) in mTeSR1 medium supplemented with 10 µM ROCKi. One day after seeding, cells were washed once with 1x Dulbecco's Phosphate-Buffered Saline (DPBS, Sigma) and medium was switched to NeuroBasal Medium/NBM [1:1 mixture of DMEM/F12, HEPES (Gibco) with Neurobasal Medium (Gibco), 0.5% (v/v) N2 supplement (Gibco), 0.5% (v/v) B27 supplement without vitamin A (Gibco), 1% (v/v) GlutaMAX (Gibco) and 10 U/mL Penicillin-Streptomycin (Gibco)] supplemented with 2 µg/mL DOX (Doxycycline Hyclate, Sigma) to induce NGN2 expression (day 0). On days 1-2, full medium changes were performed, and DOX was continuously added to the NBM medium at a concentration of 2 µg/mL. On day 3, the resulting immature NGN2-inducible neurons were dissociated with Accutase (Sigma) and seeded on 96-well µclear plates (Greiner Bio-One) or 6-well plates (Corning) coated with Poly-L-ornithine solution (PLO, Sigma) diluted 1:2 in 1x DPBS overnight followed by a mixture of 5 µg/mL of human (rhLaminin-511, Biolamina) and 10 µg/mL of mouse laminin (Sigma), at a density of 20,000 or 400,000 cells/well, respectively, in NBM medium supplemented with 10 µM ROCKi and 2 µg/mL DOX. One day later, 90% of NBM medium was changed and DOX induction at 2 µg/mL was maintained in NBM medium. On day 5, medium was fully changed to Neuronal Maturation Medium/NMM [NBM as base medium, with additional of 20 ng/mL BDNF (R&D Systems), 10 ng/mL GDNF (R&D Systems), 50 µM db-cAMP (Sigma), 200 µM L-AA2P (Sigma)], and supplemented with 2 µg/mL DOX and 10 µM DAPT (Sigma) to avoid the growth of proliferative cells. After day 5 and until completion of the experiments, half-medium changes were performed twice a week with NMM supplemented with 2 µg/mL DOX until sampling.

### Transfection of neuroblastoma cell lines and transduction of NGN2-inducible neurons

SK-N-MC cells were seeded in uncoated 96-well µclear plates (Greiner Bio-One) at a density of 20,000 cells per well and 24 hours later transfected with serially diluted ASOs (Janssen Biopharma and Axolabs GmbH) by free delivery in the culture medium, starting at 20 µM. SH-SY5Y cells were seeded in uncoated 24-well plates (Falcon) at a density of 150,000 cells per well and 24 hours later transfected with either Accell siRNAs (Dharmacon) by free delivery in the culture medium, or with a final concentration of 2 µM siRNA sequences provided by Simone et *al.*[5] (Ambion) using RNAiMax (Invitrogen) transfection reagent following manufacturer's instructions. After 48h, cells were collected for RNA and protein analysis.

NGN2-inducible neurons were transfected at DIV8 at a density of 20,000 to 25,000 cells/well with either serially diluted ASOs (Janssen Biopharma/Axolabs) or Accell siRNAs (Dharmacon) by free delivery in the culture medium. Their efficacy in knocking down *MAPT-AS1* levels was evaluated by RT-qPCR. A non-targeting ASO with a non-targeting sequence (Scrambled ASO) was used as negative control. Two ASOs were used as positive controls, one targeting *MALAT1* and another one targeting *MAPT* mRNA. Similarly, one siRNA targeting *GAPDH* and an additional one targeting *MAPT* were used as positive controls for the siRNA-based experiments.

Neurons were transduced with the same custom designed lentiviral constructs previously described for the generation of SH-SY5Y stable cell lines and with an additional construct overexpressing the entire cDNA sequence of *MAPT-AS1* (NR_024559.1) except for the PolyA-tail sequence under the cytomegalovirus (CMV) promoter (CMV:*MAPT-AS1*) at different MOI ratios. MOI = Virus Titer / Number of Cells. After 10 days of treatment, neurons were collected for RNA and protein analysis.

Information on lentiviral constructs, siRNAs and ASOs is available at Supplementary Tables S1, S2 and S3, respectively.

### RNA Isolation from Human Brain Samples

Hippocampal and entorhinal cortex tissue samples were obtained from the London Neurodegenerative Diseases Brain Bank and collected in accordance to British legislation and their ethical board[11]. The human study was evaluated and approved by the ethical committees of Leuven University and UZ Leuven[11]. Total RNA from the human brain tissue was extracted by homogenization in TRIzol (Invitrogen) using 1 ml syringes and 22G/26G needles and purified on mirVana spin columns according to the manufacturer's instructions (Ambion). RNA purity (260/280 and 260/230 ratios) and integrity were assessed using Nanodrop ND-1000 (Nanodrop Technologies) and Agilent 2100 Bioanalyzer with High Sensitivity chips (Agilent Technologies, Inc.) and Qubit 3.0 Fluorometer (Life Technologies), respectively. A total of 300 ng of RNA was used per sample for cDNA conversion and RNA levels evaluated by RT-qPCR. Range of Ct values for each of the transcripts were as follows: no amplification for *t-NAT2l*; 30.9-31.2 for *t-NAT2s*; 34.8-35.2 for *t-NAT1*; 30.5-31.2 for total *t-NAT*; 36.3-37.4 for *MAPT-AS1* (Taqman).

### Multiplex fluorescent *in situ* hybridization (RNAscope) in Human Brain Samples

Human brain samples were obtained from the Netherlands Brain Bank (NBB), Netherlands Institute for Neuroscience, Amsterdam. Written informed consent was given by the donors for brain autopsy and for the use of material and clinical data for research purposes, in compliance with national ethical guidelines[12]. Frozen human brain blocks of superior frontal gyrus from 6 individuals were cryosectioned to a thickness of 10µm using a CryoStar NX70 cryostat (ThermoFisher), layered onto SuperFrost Plus glass slides (ThermoFisher) and further stored at -80° C before experiments. Sectioned samples on glass slides were processed for in situ hybridization, which was performed using the RNAscope Multiplex Fluorescent V2 Assay (ACD Bio-Techne) according to the manufacturer's instructions. Sections were fixed in cold 4% PFA and dehydrated using a series of ethanol dilution steps, followed by treatment with Hydrogen Peroxide for 10 minutes at RT. Protease digestion using Protease IV provided in the RNAscope kit was carried out for 20 minutes at RT. Hybridization proceeded for 2 hours at 40°C. The following ACD Bio probes were used: Hs-*RBFOX3*-C1 (415591), Hs-*MAPT-AS1*-C2 (564491-C2), and Hs-*MAPT*-C3 (408991-C3). Hs-*POLR2A*-C1; *PPIB*-C2; *UCB*-C3 were used low-, medium- and high-expressing positive control probes, respectively. Bacterial *DapB* probe was used as a negative control. Slides were stored in 5X SSC buffer overnight at RT. Amplification was carried out the following day using the three amplification reagents provided with the kit. Detection was done with TSA® Plus Fluorophores FITC (*MAPT-AS1*), Cy5 (*MAPT*) and Cy3 (*RBFOX3*). Slides were incubated for 30 seconds with TrueBlack 1x solution and washed with 1x DPBS followed by DAPI staining. Images were acquired using the Leica SP8× confocal microscope and analysed using the ImageJ software.

### RNA isolation from SK-N-MC, SH-SY5Y and NGN2-inducible neurons

Total RNA from SK-N-MC, SH-SY5Y and NGN2-inducible neurons was extracted using the RNeasy 96 kit or RNeasy Mini Kit (Qiagen) according to manufacturer's recommendations with minor changes. Briefly, cell culture medium was removed and RLT lysis buffer (Qiagen) was added to the cells for 20 minutes at RT with shaking at 500 rpm. No β-mercaptoethanol was added to the RLT lysis buffer. Lysates were either stored at -80°C and thawed prior to RNA isolation or protocol was carried out immediately as described in the kits. On-column DNase treatment was performed as instructed by the manufacturer. After RNA elution with RNAase-free water, RNeasy columns/ column plates were discarded and the eluate tubes/ plates containing the total RNA were stored at -80°C until used for cDNA conversion. RNA quantification was performed using the absorbance 260nm value with NanoDrop.

### RNA Subcellular Fractionation in NGN2-inducible neurons

RNA fractionation was performed using the RNA Subcellular Isolation kit (Active Motif) according to the manufacturer's instructions. Briefly, NGN2-inducible neurons were washed with 1x DPBS dissociated from 6-well plates with Accutase (Sigma). Then, NBM medium was added to each well to detach the cells from the plate and cells were centrifuged at 450 x g for 5 minutes. Cells were washed with 1x DPBS and centrifuged at 14,000 rpm for 5 minutes. To isolate cytoplasmic and nuclear fractions, each pellet was resuspended in 120 µl of Complete Lysis Buffer and incubated on ice for 10 minutes. Then, cells were centrifuged at 14,000 rpm for 5 minutes at 4°C. Supernatant (cytoplasmic fraction) was transferred to a new tube without disturbing the pellet (nuclear fraction). Complete Buffer G was added to each fraction as recommended. To extract the RNA fractions, lysates were mixed with 1 volume of 70% ethanol by pipetting up and down 3 times and transferred to the corresponding purification columns. To promote RNA binding to the silica membrane, columns were centrifuged at 14,000 rpm for 1 minute at 4°C. The eluate was discarded, and columns were washed with Wash Buffer by centrifugation at 14,000 rpm for 1 minute at 4°C. The eluate was discarded, and the columns were washed with 70% Ethanol using the same centrifugation parameters. The eluate was discarded, and a final centrifugation step was performed at 14,000 rpm for 2 minutes at 4°C. Then, purification columns were transferred to a new collection tube and 50 µl of RNAse-free water was added to each column. To elute the RNA, columns were centrifuged at 14,000 rpm for 1 minute at 4°C. Purification columns were discarded and the eluate tubes containing the

RNA subcellular fractions were stored at -80°C until used for cDNA conversion. RNA quantification was performed using the absorbance 260nm value with NanoDrop.

### cDNA conversion and Real-Time quantitative PCR (RT-qPCR)

Conversion of total RNA to first-strand cDNA was carried out using the High-Capacity cDNA conversion kit in a 20 µl reaction following manufacturer's instructions (ThermoFisher) without RNAse inhibitor in the 2x RT master mix reaction. cDNA samples were stored at -20°C until analyzed with RT-qPCR.

RT-qPCR experiments were performed in a 10 µl reaction and run for 40 cycles. PrimeTime Gene Expression Master Mix (IDT DNA) or PowerUp SYBR Green Master Mix (ThermoFisher) were used with the appropriate Taqman assays or primers, respectively. RT-qPCR reactions were run in triplicates. RT-qPCR was performed with the QuantStudio™ 7 Flex Real-Time PCR System (Design & Analysis Software v2.3.0; ThermoFisher) or QuantStudio 12K Flex Real-Time PCR System (QuantStudio Flex Software v1.2.4; ThermoFisher). Results were analyzed using qBase+ software v3.2 (Biogazelle)[13]. For each experiment, 8 different reference genes were run, and their stability was determined using GeNorm analysis[13]. Target gene expression levels were normalized to the 2 most stable reference genes and calibrated to a control sample/group unless specified otherwise. All expression levels are shown as calibrated normalized relative quantities (CNRQ)[13] unless specified otherwise. Information on custom designed primers and IDT DNA pre-designed Taqman RT-qPCR assays and primers is available at Supplementary Table S4.

### Protein Isolation and Tau detection with Mesoscale Discovery (MSD)

Protein samples were obtained by lysis in cold RIPA buffer (Sigma) supplemented with phosphatase (PhosStop, Roche) and protease inhibitors (cOmplete, Roche). Lysates were stored at -20°C prior to the protein analysis.

To quantify Tau protein levels using Mesoscale (MSD) platform, JRD/hTau/24 (same as hTau21, Janssen R&D)[14] capture antibody was diluted in 1x DPBS at a final concentration of 1 µg/mL and added directly into the wells on 96-well sector standard plates (L15XA, MSD). Plates were then sealed and incubated overnight at 4°C. After overnight incubation, coated plates were blocked with 0.1% (v/v) Blocker Casein (ThermoFisher) in 1x DPBS for 1-2 hours at RT with agitation at 500 rpm. Following this step, plates were washed 5 times with washing buffer [0.05% Tween (v/v) in 1x DPBS]. After washing, standards and samples were added to the plates diluted in RIPA buffer and incubated overnight at 4°C with shaking at 500 rpm.

Human recombinant tau441 protein (TAU-4R-WT, Tebu Bio) was used to generate standard curves and to interpolate Tau protein concentrations. The next day, plates were washed again 5 times as previously described, and the JRD/hTau/43 (Janssen R&D)[15] detection antibody was added at a 1:3000 dilution to the plates and incubated for 2 hours at RT with agitation at 500 rpm. After this step, plates were washed as before, and MSD Read Buffer T (MSD) with surfactant diluted 1:2 in distilled water was added to each well. Plates were immediately read using the MSD SECTOR Imager 6000 (MSD). Tau concentration (ng/mL) was interpolated from human recombinant 2N4R Tau standard curve.

### Western Blot analysis

SH-SY5Y protein samples were obtained by lysis in cold RIPA buffer (Sigma) supplemented with phosphatase (PhosStop, Roche) and protease inhibitors (cOmplete, Roche). Lysates were stored at -20°C prior to the protein analysis. Protein lysates concentrations were measured using a BCA Protein Assay Kit (ThermoFisher). Proteins were separated in 4–12% SDS–polyacrylamide gel (Criterion XT Bis-Tris, Bio-Rad) in MOPS buffer and transferred to 0.2-μm nitrocellulose membrane for 7 min at 2.5A constant (pre-defined Midi gel mixed molecular weight protocol), using the Trans-Blot Turbo Transfer system (Bio-Rad). Immunoblotting of SH-SY5Y was performed with the following primary antibodies: DAKO Polyclonal rabbit anti-human Tau (1:15000; Agilent #A0024) and monoclonal mouse anti-β-actin (1:2000; Sigma #A2228). Secondary antibodies were as follows: donkey-anti-rabbit IgG-HRPO (1:5000; GE Healthcare #NA934V) and goat-anti-mouse IgG-HRPO (1:3000; BioRad #170-6516). Membranes were developed with Amersham Imager 600 (GE Healthcare) and quantified using ImageJ-Fiji (version v1.53c).

### Immunofluorescence

Medium was carefully removed from NGN2-inducible neurons and cells were fixed with 4% (v/v) paraformaldehyde (ThermoFisher) for 15 minutes at RT. Then, neurons were blocked with 5% (v/v) normal goat serum (Sigma) diluted in 1x DPBS supplemented with 0.3% (v/v) Triton X-100 (ThermoFisher) for 1 hour at RT, followed by overnight incubation at 4˚C with primary antibodies against Oct3/4 (1:50; SantaCruz #sc-5279), Sox2 (1:250; Cell Signalling #3579S), Ngn2 (1:500; Millipore #AB5682), Nestin (1:250; Millipore #MAB5326), NeuN (1:1000; Millipore #ABN78), Tuj1 (1:1000; Abcam #AB52623), HT7 (1:250; ThermoFisher MN1000) and Map2 (1:7000; Abcam #AB5392). The following day, cells were washed 3 times for 5 minutes in 1x DPBS, and then incubated for 60 minutes with Alexa-Fluor conjugated secondary antibodies (ThermoFisher: Mouse AF488, #A-11029; Rabbit AF568, #A-11011; Chicken AF647, #A-21449) diluted 1:400 in 1x DPBS supplemented with 0.3% (v/v) Triton X-100 at RT. Images were acquired using the Opera Phenix Plus High Content Screening System confocal microscope.

### Antisense oligonucleotide synthesis

Antisense oligonucleotides (ASOs) were synthesized as previously described[16] by Janssen Biopharma (South San Francisco, USA) or by Axolabs GmbH (Germany). All ASOs are 20 nucleotides long. The 5 nucleotides on either end contain 2'-O-methoxyethyl (2'MOE) nucleotides whereas the central segment comprises 2'-deoxynucleotides. All cytosine residues are 5'-methylcytosines. All inter-nucleotide linkages are phosphorothioate. The lyophilised ASOs are reconstituted to a stock concentration of 1 mM in 1x DPBS. The *MAPT-AS1* ASOs tiling the predicted mature transcript (NR_024559.1, 840 bp long) with no overlaps were designed without any restrictions.

### Statistical analysis

All data are presented as mean and standard deviation (SD) unless specified otherwise. Statistical significance was set at $\alpha=0.05$. Statistical analysis was performed using Graphpad Prism (v8.4.2). Normality analysis was always performed prior to choosing the post-hoc test to analyse the data. Statistical analysis of expression data from human brain samples was performed using parametric unpaired t-test for normally distributed groups or nonparametric Mann-Whitney U-test for non-normal data. For multiple comparison datasets, One-Way ANOVA with Dunn's multiple comparisons test was used, unless specified otherwise. More details on each individual statistical analyses can be found in the relevant figure legends.
