## Supplementary Tables for "The MIR-NAT *MAPT-AS1* does not regulate Tau expression in human neurons"

*Table S1. Customized VectorBuilder lentiviral constructs information. VectorBuilder Virus ID, Abbreviation, Target RNA/Protein, Viral Type, Promotor and Titer are shown.*
*Abbreviation corresponds to the name given in this manuscript.*

| Virus ID | Abbreviation | Target RNA/Protein | Viral type | Promoter | Titer (1st batch) | Titer (2nd batch) |
|---|---|---|---|---|---|---|
| VB190716-1013rtr | CMV:MAPT-AS1 | MAPT-AS1 without polyA | Lentivirus | CMV | 2.73 x 10e8tu/mL | |
| VB190716-1208fkr | SYN1:MAPT-AS1 | MAPT-AS1 without polyA | Lentivirus | SYN1 | 4.03 x 10e8tu/mL | 1.17 x 10e9tu/mL |
| VB170504-1049ajc | SYN1:eGFP | EGFP | Lentivirus | SYN1 | 7.5 x 10e8tu/mL | 1.04 x 10e9tu/mL |
| VB180802-1002wky | CMV:eGFP | EGFP | Lentivirus | CMV | 8.1 x 10e8tu/mL | |
| VB210531-1081cph | SYN1:miniNAT | miniNAT (Simone et *al.* ) | Lentivirus | SYN1 | 4.12 x 10e8tu/mL | |

*Table S2. Customized and commercially available Accel siRNAs sequences (Dharmacon) and pre-designed and custom-designed Silencer Select siRNAs (Ambion; Simone et. al publication).*
*Full Name, Abbreviation, Reference, Sense and Antisense Sequences are shown.*
*Abbreviation corresponds to the name given in this manuscript.*

| Full Name | Abbreviation | Reference | Sense Sequence (5'-3') | Antisense Sequence (5'-3') |
|---|---|---|---|---|
| Accel MAPT-AS1 s237111 | MAPT-AS1 siRNA | HR1ZN-008811 | CCACUUCAUGGAUAAGUAAUU | UUACUUAUCCAUGAAGUGGUU |
| Accell GAPD siRNA | GAPDH siRNA | D-001930-01-50 | AATTACTTATCCATGTTGTGG | AACCACTTCATGGATAAGTAA |
| Accell Human MAPT (4137) siRNA | MAPT siRNA | A-012488-13 | | |
| | | A-012488-14 | | |
| | | A-012488-15 | | |
| | | A-012488-16 | | |
| Accell Non-targeting Control Pool | Scrambled siRNA | D-001910-10-20 | | |
| Ambion siNT1nover | NT1 siRNA | 4390828 | CGGCGAGGCAGAUUUCGGAtt | UCCGAAAUCUGCCUCGCCGtc |
| Ambion siNT2nover | NT2 siRNA | 4390828 | GCCGCCGAGUCCGUCCACAtt | UGUGGACGGACUCGGCGGCcg |
| Ambion siEx4-n268298 | Exon4_1 siRNA | 4390815 | GAUUUGUCAUGAGUCUCUUtt | AAGAGACUCAUGACAAAUCaa |
| Ambion siEx4-n268302 | Exon4_2 siRNA | 4390815 | AGGACAAUGUCCUAAGGAAtt | UUCCUUAGGACAUUGUCCUcc |
| Ambion Negative Control #2 siRNA* | Scrambled siRNA | 4390847 | | |

** SH-SY5Y experiment only*

*Table S3. ASOs sequences. ASO name, Target Gene, Sequence, Wing Chemistry and Gapmer Chemistry are shown.*
*ASO name corresponds to the name given in this manuscript.*

| ASO name | Target Gene | Sequence (5'- 3') | Wing chemistry | Gapmer chem |
|---|---|---|---|---|
| MAPT-AS1 ASO10 | Human MAPT-AS1 | AAGATCATGTCTCTCTCTTG | 2'MOE OPS | OPS |
| MAPT-AS1 ASO16 | Human MAPT-AS1 | CTTTGCTGTGTCAATGTGGG | 2'MOE OPS | OPS |
| MAPT ASO | Human MAPT | Undisclosed | | |
| Scrambled ASO | Non-Targeting | CCTTCCCTGAAGGTTCCTCC | 2'MOE OPS | OPS |
| MALAT1 ASO | Human MAPT-AS1 | UGCCUTTAGGATTCTAGACA | 2'MOE OPS | OPS |

*Table S4. GeNorm human reference genes, custom designed primers and IDT DNA pre-designed Taqman RT-qPCR assays and primers.*

*GeNorm human reference genes; Gene name, Primer and Sequence are shown. FWD = forward; REV = reverse*

| **Gene Name** | **Primer** | **Sequence** |
|---|---|---|
| GAPDH | FWD | AAGGTGAAGGTCGGAGTCAAC |
| | REV | GGGGTCATTGATGGCAACAATA |
| RNF20 | FWD | TTATCCCGGAAGCTAAACAGTGG |
| | REV | GTAGCCTCATATTCTCCTGTGC |
| VIPAR | FWD | GGGAGACCCAAAGGGGAGTAT |
| | REV | GGAGCGGAATCTCTCTAGTGAG |
| SCLY | FWD | ACTATAATGCAACGACTCCCCT |
| | REV | CTTCCTGCTGAATACGGGCTG |
| PRDM4 | FWD | CACCTCCACAGTACATCCACC |
| | REV | TGATAGGGATCTAGTGCTGAAGG |
| ENOX2 | FWD | TCATTGTGGAAGTTTTCGAGCA |
| | REV | TGCGGTAACCAGACAGATACA |
| UBE4A | FWD | TAGCCGCTCATTCCGATCAC |
| | REV | GGGATGCCATTCCCGCTTT |
| UBE2DE | FWD | CAGTCCCTATCAGGGTGGAGT |
| | REV | AAGGGGTAATCTGTTGGGAAATG |
| ERCC6 | FWD | TCACGTCATGTACGACATCCC |
| | REV | GTGGCAGCTTGAGGGCTAAG |

*Custom designed primers; Gene name, Primer and Sequence are shown. FWD = forward; REV = reverse*

| **Gene Name** | **Primer** | **Sequence** |
|---|---|---|
| MAPT (Assay 1) | FWD | CCTCCAAGTGTGGCTCATTA |
| | REV | CAATCTTCGACTGGACTCTG |
| MAPT (Assay 2) | FWD | CAGTGGTCCGTACTCCA |
| | REV | TGGACTTGACATTCTTCAGG |
| 3R MAPT | FWD | AGGCGGGAAGGTGCAAATA |
| | REV | GCCACCTCCTGGTTTATGATG |
| 4R MAPT | FWD | CGGGAAGGTGCAGATAATTAA |
| | REV | TATTTGCACACTGCCGCCT |
| MAPT (Simone et *al.* ) | FWD | GATTGGGTCCCTGGACAATA |
| | REV | GTGGTCTGTCTTGGCTTTGG |
| Total t-NAT (Simone et *al.* ) | FWD | GGAGTCAGAACAAGGACGGG |
| | REV | GCACATCCTGGGCTACTGTT |
| t-NAT2l (Simone et *al.* ) | FWD | CCAAGACTCCAGTTCTCGCC |
| | REV | CATCCTGGGCTACTGTTCCA |
| t-NAT2s (Simone et *al.* ) | FWD | ACCTCCTGTCCAGGCTTCT |
| | REV | CCGCACACTAACTGCTTTGA |
| t-NAT1 (Simone et *al.* ) | FWD | GAGGAGGAGAAGGTGGCTGT |
| | REV | GGACCTGGTTCCCTTCACTT |
| miniNAT (Simone et *al.* ) | FWD | CGGGAGAGGTTAATACACCCA |
| | REV | CTGTGTCAATGTGGGCTTCT |

* used for Figure 1i-j (*MAPT* mRNA levels shown as average of Assays 1 and 2).

*IDT DNA pre-designed Taqman RT-qPCR assays and primers (Available at https://eu.idtdna.com/site/order/qpcr/predesignedassay).*
*Gene name, Assay ID, and Assay Configuration are shown.*

| **Gene Name** | **Assay ID (IDT DNA)** | **Assay Configuration** |
|---|---|---|
| MAPT-AS1 (Assay 1) | Hs.PT.58.20559679 | Std, FAM/ZEN/IBFQ, P:P 2 |
| MAPT (Assay 3)* | Hs.PT.58.28269192 | Std, DNA Primer |
| GFAP | Hs.PT.58.1057167 | Std, DNA Primer |
| TREM2 | Hs.PT.58.40294042 | Std, DNA Primer |
| RBFOX3 | Hs.PT.58.2776427 | Std, FAM/ZEN/IBFQ, P:P 2 |
| TUBB3 | Hs.PT.58.20385221 | Std, DNA Primer |
| MALAT1 | Hs.PT.58.26451167.g | Std, FAM/ZEN/IBFQ, P:P 2 |

* used for Extended Data Figure 1j.
